# Epigenetic gene regulation is controlled by distinct regulatory complexes utilizing specialized paralogs of TELOMERE REPEAT BINDING FACTORS

**DOI:** 10.1101/2025.11.10.687574

**Authors:** Maik Mendler, Kristin Krause, Simone Zündorf, Prathamesh Sannak, Petra Tänzler, Sara Stolze, Hirofumi Nakagami, Franziska Turck

## Abstract

Epigenetic regulators shape chromatin landscapes, allowing cells to express distinct gene sets depending on cell-type, developmental stage or environmental cues. These regulatory complexes rely on interactions with sequence-specific DNA binding proteins, such as the small family of TELOMERE REPEAT BINDING FACTORS (TRBs). TRBs are components of chromatin regulatory complexes with opposing functions, such as the epigenetic repressors Polycomb Repressive Complex 2 (PRC2) and a JMJ14/NAC complex that respectively add and removes the repressive H3K27me3 and positive H3K4me3 modification, but also with the plant-specific PEAT complex that is linked to histone acetylation and gene activation.

We dissected the partial redundancy between TRB1, TRB2 and TRB3 in target gene selection and interaction with different chromatin regulatory complexes. High redundancy of TRBs is suggested by major phenotypic changes that are only observed *trb* triple mutants; however, we found different target site preference between TRB1-3 and preferred partnership with chromatin complexes. Furthermore, TRB paralogs interacted with the NuA4 histone acetylation complex, both together with and in absence of PEAT. Among the three paralogs, TRB1 had more unique binding sites and correlated stronger with PEAT and NuA4 functions. In contrast, TRB2 and TRB3 were more dependent on the presence of *bona fide* telo-box motifs and were more likely to be found at PRC2 associated sites. Overall, we provide insight into the diverse roles of TRBs in epigenetic gene regulation and how their diversification contributes to their apparent redundancy, as well as their observed activating and repressing effects on gene expression.

## Background

In *Arabidopsis thaliana*, telo-box motifs are widely found at gene regulatory elements; furthermore, they are native to the telomeres at chromosomal ends, where they occur as direct repeats (TTTAGGG × n; n □ 2 to 1000+) and associate with several telomere repeat binding proteins, including TELOMERE REPEAT BINDING FACTOR (TRB) 1-5 [1–3]. TRB proteins belong to the Single myb histone (Smh) family and contain an N-terminal myb, a H1/H5-linker and a C-terminal coiled-coil domain [4]. The direct interaction of TRBs with telomere repeats is mediated by their N-terminal myb-domain, which belongs to the telobox class of myb-domains that is shared among telomere repeat binding proteins in all eukaryotes [5, 6]. TRBs directly interact with the telomerase subunit TERT and are thought to be part of the plant shelterin complex which aids the telomerase in solving the end-replication problem and protects telomere ends from being falsely recognized as DNA double strand breaks [1, 7]. The H1/H5-linker histone domain is involved in the formation of TRB multimers at telomere ends. For TRB1, it was shown that binding to interstitial telomeric repeats, which are relics of chromosomal fusions specific to *A. thaliana*, is prevented by the presence of linker histone H1, indicating that while the H1/H5 domain may contribute to target binding, it cannot outcompete the canonical linker histone [8]. The C-terminal coiled-coil domain is thought to mediate the interaction with other proteins [5, 9]. Telo-box motifs were first linked to promoter regions of highly expressed genes of the translation machinery [3]. Genome-wide binding analysis of TRB1 confirmed the link to genes encoding for the translational apparatus [9, 10].

TRB1 is an integral component of the plant-specific PWWP-ENHANCER OF POLYCOMB-LIKE-ARID-TRB (PEAT) complex, which is predominantly involved in gene activation [11, 12]. PEAT activates target genes through a dual approach that involves histone acetylation and deubiquitination of mono-ubiquitinated H2A (H2AKub1) [11]. Histone acetylation is facilitated through HISTONE ACETYLASE RELATED TO MYST (HAM) 1 and 2 which links PEAT to the Nucleosomal Acetyltransferase of histone H4 (NuA4) complex, which co-purifies with PEAT components [11]. Co-purification of UBIQUITIN SPECIFIC PROTEASE 5 (UBP5) and PEAT positions PEAT as a direct antagonist of the epigenetic repressor Polycomb Repressive Complex (PRC) 1, which sets the H2AKub1 mark [13].

Mutated alleles of *TRB1* and *TRB3* were identified as genetic enhancers of mutations in *LIKE-HETEROCHROMATIN PROTEIN 1* (*LHP1*) and *CURLY LEAF* (*CLF*). LHP1 and CLF act as accessory and integral part of PRC2, respectively [9, 14] and LHP1 also interacts with PRC1 components that catalyze H2AKub1 [15]. PRC2 establishes the covalent modification tri-methylation of histone H3 at lysine 27 (H3K27me3) at thousands of loci, resulting in transcriptional repression of target genes (reviewed by [16, 17]). TRBs can recruit PRC2 to telo-box motifs via their direct interactions with the PRC2 components CLF/SWN, which is essential for stable H3K27me3 coverage and epigenetic repression of a subset of these genes [14]. While TRB1 binding sites were under-represented within H3K27me3 marked regions, telo-box motifs were overrepresented, in particular at regions that show reduced H3K27me3 in *clf* mutants [18]. Telo-box motifs were also enriched in regions bound by the PRC2 components FERTILISATION INDEPENDENT ENDSPERM (FIE), SWINGER (SWN) and CLF [19–21]. Finally, TRB1-3 are also part of a transcription repressive complex composed of the histone de-methylase JUMONJI (JMJ14), NAC50 and NAC52 (names after the founding family members No Apical Meristem (NAM1), ARABIDOPSIS TRANSCRIPTION FACTOR (ATAF1/2), CUP SHAPED COTYLEDONE (CUC2)) [22, 23].

The association of telo-box motifs and TRBs with repressed as well as highly expressed genes and with repressive as well as activating chromatin complexes raises the question of how the functional context is established and to which extent TRBs are specialized in their molecular function. Phenotypic analysis confirmed the high redundancy between TRB1-3, as strong phenotypic changes were only observed in triple mutants. In contrast, comparison of genome-wide binding revealed many sites exclusively bound by TRB1 that were more likely associated with PEAT/NuA4 or NuA4. In contrast, binding sites preferred by TRB2 and TRB3 over TRB1 were the most highly associated with PRC2-mediated gene repression.

## Results

### Transcriptomic changes in *trb* double and triple mutants indicate a limited partial redundancy between paralogs

As we and others have previously reported, plants homozygous for two mutated alleles of *TRB1*, *TRB2* or *TRB3* are indistinguishable from wild-type controls grown under standard conditions, but a deletion of the third allele has a catastrophic effect on plant development [14, 23]. To evaluate the extent of genetic redundancy, we grew plants that still segregated one functional *TRB* allele (*prope triple*) together with all double mutant combinations and Col-0 controls in standard growth conditions (LD, 16h light/8h dark, 21°C). Triple *trb123* mutants were strongly dwarfed and usually died before their *prope triple* or double homozygous siblings flowered (Figure 1A). While *prope triple* mutants that segregated functional alleles of *TRB1* and *TRB3* flowered as controls, plants carrying only one functional allele of *TRB2* flowered significantly earlier indicating that the latter is a slightly weaker paralog with respect to a role in flowering time regulation (Figure 1B).

**Figure 1.**
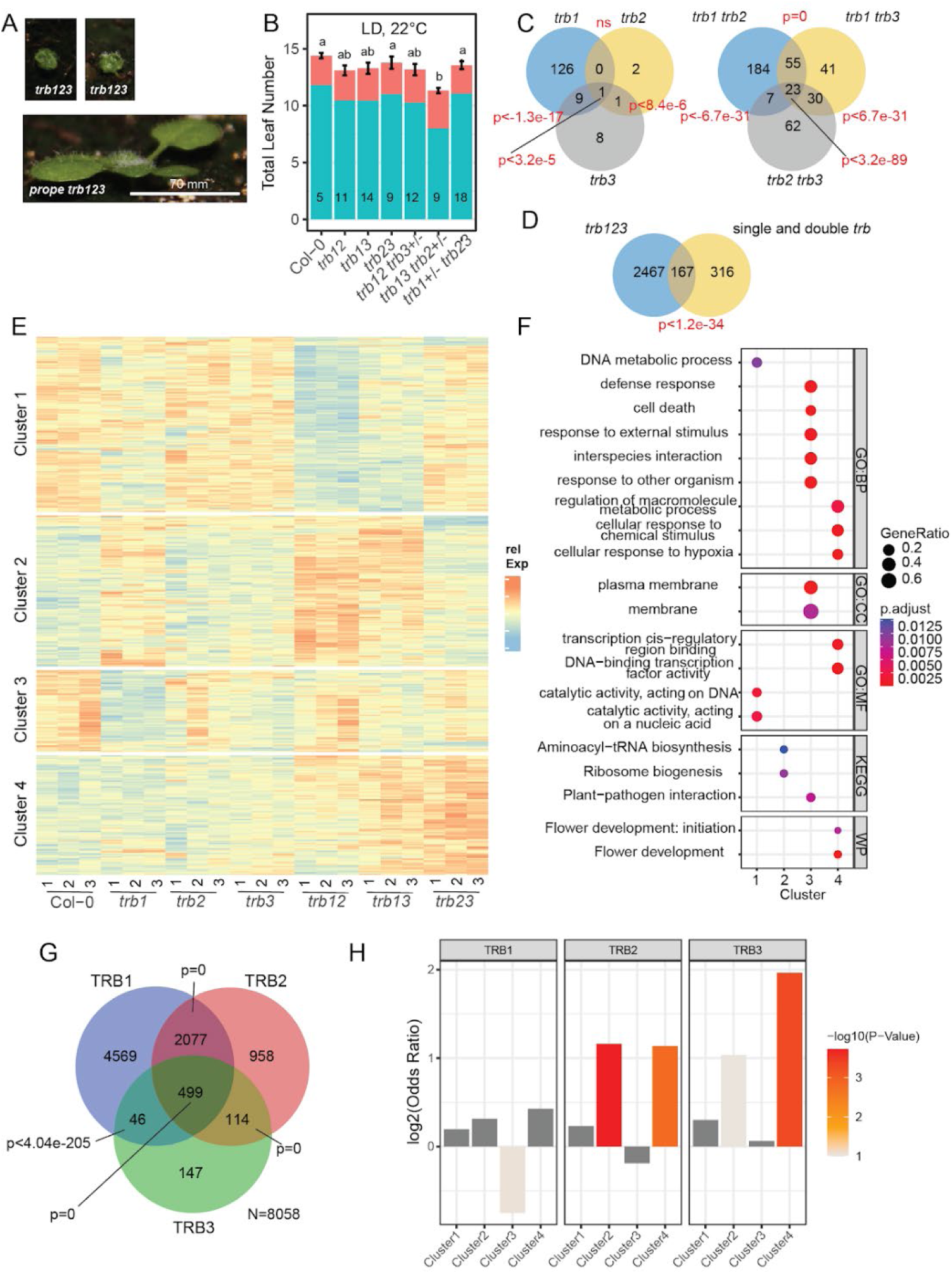
Limited partial redundancy of TRB1-3. **A)** Phenotypes of trb123 triple mutants and siblings with one functional TRB allele. Common scale bar indicated. **B)** Flowering time of double trb mutant and prope triple mutants scored as total leaves on main shoot (rosette leaves: green, cauline leaves: red). Plants were grown at 22°C in long days (16h light/8h dark) in culture chambers. Significant differences determined by ANOVA, letters indicate HSD groups at p<0.01. Replicates (5 < n < 18) are indicated within bars. Error bars show standard error of the mean. **C)** Venn diagrams showing the number of differentially expressed genes (DEGs) in single trb (left) and double trb mutants (right) compared to Col-0. **D)** as C) comparing DEGs in triple trb123 mutants against all DEGs in C). Significance of overlap in C) and D) determined by SuperExact test using all expressed genes as background. **E)** Heatmap of clusters of all DEGs as in C). Clustering was performed after Variance Stabilization Transformation (VST) of read count data and normalization by the mean for each gene. **F)** Gene-Onthology (GO) and pathway enrichment analysis for clusters shown in E). Significance was tested against the background of all expressed genes. Relative enrichment of GO terms for Biological processes (GO:BP), cellular compartment (GO:CC), molecular function (GO:MF) and KEGG metabolic pathways and wikipathways (WP) are indicated by bubble sizes, statistical significance by color codes. **G)** Venn diagram showing the number of genes associated with ChIP-seq peaks identified for TRB1, TRB2 and TRB3. Statistical test as in C using all annotated genes as background. **H)** Distribution of TRB1, TRB2 and TRB3 target genes among the transcriptional clusters shown in E). Bar plots show Odds ratio and −log10 pValues as determined by Fisher’s Exact test are indicated as color code.

Both, high redundancy and signs of unequal redundancy, were also observed at the level of transcriptome changes. Using above ground tissue of 14-day-old seedlings grown on soil in standard growth conditions (LD, 16h light/8h dark, 21°C), we found overall 483 differentially expressed genes (DEGs) in *trb* single and double mutants compared to Col-0 (Figure 1C and Supplemental File 1). Of these DEGs, most were specific to *trb1* single and *trb1 trb2* double mutants, indicating that *TRB2* and *TRB3* cannot fully compensate for the loss of *TRB1*. Only around a third (35%) of the DEGs identified in single and double mutants were shared with *trb123*, which represent only 6% of the 2634 DEGs identified in these mutants (Figure 1D). While a higher number of DEGs is expected from the strong phenotypic changes observed in *trb123*, the occurrence of genes specific to the milder mutants is more difficult to rationalize.

To gain more insight on the specific impact of *TRB* paralogs on the transcriptome, we performed partitioning around medoid (PAM)-clustering of all mis-regulated genes based on their relative expression profile. The number of clusters k=4 was empirically determined as best (Figure 1E). Cluster 1 and 2 correlated with the loss of *TRB1* function, showing respectively decreased- and increased expression. GO-term enrichment analysis associated cluster 1 with functions in DNA damage and repair, while cluster 2 enriched GO-terms related to ribosome biogenesis (Figure 1F, Supplemental File 1). Cluster 3 showed decreased expression in *trb1*, *trb3* and *trb13* double mutants and enriched GO-terms related to responses to biotic stress. Cluster 4 was mostly determined by the combination of *trb3* with either *trb1* or *trb2*, showing increased expression of genes in these double mutants. Cluster 4 enriched the wiki-pathway flowering initiation and flowering development (https://www.wikipathways.org/) based on the presence of the floral organ identity genes *AGAMOUS* (*AG*), *SEPELLATA* (*SEP*) *1* and *3*, the flowering time regulator *TEMPRANILLO 1* (*TEM1*) and the PcG component *EMBRYONIC FLOWER 1* (*EMF1*), which were previously identified as TRB-dependent targets of PRC2 [14] (Supplemental Figure 1).

### Binding sites of TRB1-3 reveal distinct target site preferences

To better understand the redundant and specific roles of TRB paralogs, genome-wide binding profiles of all three were compared. TRB ChIP-seq libraries were prepared using chromatin from 14-day old seedlings stably expressing *TRB2* and *TRB3* carboxy-terminal fusions to *YELLOW FLUORESCENT PROTEIN* (*YFP*) under the control of their respective promoters (*TRB2-YFP* in Col-0 background and *TRB3-YFP* in *trb3-2* background, [14]). TRB1-GFP ChIP-seq data were taken from Zhou *et al.* (2016). Enriched peaks were called against control ChIP-seq libraries prepared from Col-0 chromatin precipitated with the same anti-GFP antibody from two biological replicates using the Irreproducible Discovery Correction (IDR) framework at a cut-off of IDR ≤0.05 (Supplemental Figure 2,[24]). The analysis identified 7483, 3771, 845 peaks for TRB1, TRB2, TRB3 respectively (Supplemental Figure 3A and Supplemental File 2). The observed proportion of peaks for TRB paralogs is similar to those obtained in a recent study by Wang *et al.* 2023 (Supplemental Figure 3B). Visual inspection of gbrowse tracks showed that many sites unique to TRB1 showed clearly distinct peaks, while sites unique to defined as TRB2 or TRB3 frequently showed low enrichment (below the significance threshold) of the other TRBs (Supplemental Figure 3C). We annotated all TRB peaks to target genes to estimate whether transcriptional patterns detected in the double and triple mutants could be related to the binding of TRB paralogs (Figure 1G, Supplemental File 3). Overall, DEGs were slightly more likely to be TRB target genes than expected by chance (Fisher’s Test, p< 0.017, odds ratio 1.27). Next, we tested for specific TRB paralog enrichment between the four PAM-clusters. For TRB1, a direct effect on gene expression was suspected for transcriptional cluster 1 and 2 that show obvious changes in *trb1* and *trb1 trb2* mutants; however, a specific overrepresentation of TRB1 targets could not be confirmed for any of the clusters, while a mild underrepresentation was detected for cluster 3 (Fisher’s test, p<8.8×10^-2^, Figure 1H). In contrast, cluster 2 was overrepresented for TRB2 and TRB3 target genes (Fisher’s test, p< 1.8×10^-4^ and 9.2×10^-2^, respectively). While TRB2 was more significantly overrepresented in cluster 2, TRB3 was more significantly overrepresented in cluster 4 compared to TRB2 (4.8×10^-4^ and 1.5×10^-3^, respectively). Taken together, despite a large redundancy, TRB1-3 show target site preference, which is reflected in the expression patterns of a small set of genes in double or, in the case of *trb1*, single mutants. Overall, a direct more specialized role of TRB2 and TRB3 in gene repression is supported by the overrepresentation of direct targets among clusters showing increased expression in double mutants.

### TRB paralogs show preferred association with distinct chromatin regulatory complexes

To better understand the redundant and specific roles of TRB paralogs within the context of associated chromatin regulatory complexes, we compared the binding of TRB1-3 with available data for TRB-interacting protein complexes. Representative of the PEAT complex, genome-wide binding data were available for core-components EPCR1 and PWWP1 [12] and for UPB5, which removes the PRC1-associated mono-ubiquitination of H2A [13] and was recently proposed to be a PEAT component [11]. HAM1 and EPL1B are members of the NuA4-complex, the former but not the latter was also shown to interact with PEAT [11]. CLF and SWN are core components of the PRC2 complex [21]. The H3K4me3 de-methylase JMJ14 was shown to interact with TRB with a role in balancing gene expression [23]. Since data were generated by different groups and analyzed through different bioinformatic pipelines, we classified peaks into decile bins based on the published significance scores for each protein. We divided the genome of *Arabidopsis thaliana* into 238296 bins of 500 bp length and considered proteins to co-localize if any part of their associated peaks was found in the same bin (Supplemental File 3). Pearson’s correlation coefficient analysis identified four complex groups with strong correlations within the groups, corresponding to PEAT (EWPCR1, PWWP1, UBI5), NuA4 (HAM, EPL1B), TRBs-JMJ14 (TRB1, TRB2, TRB3, JMJ14) and PRC2 (CLF, SWN) (Figure 2A). Obvious positive correlation was also observed between PEAT, NuA4, TRB1 and TRB2 and between TRB3 and PRC2. In contrast, JMJ14 showed negative correlation with PEAT and NuA4 and was rather neutral with respect to PRC2 binding sites, indicating that although JMJ14 and TRBs often bind targets together, they do not usually also bind at PEAT and NuA4 associated sites.

**Figure 2.**
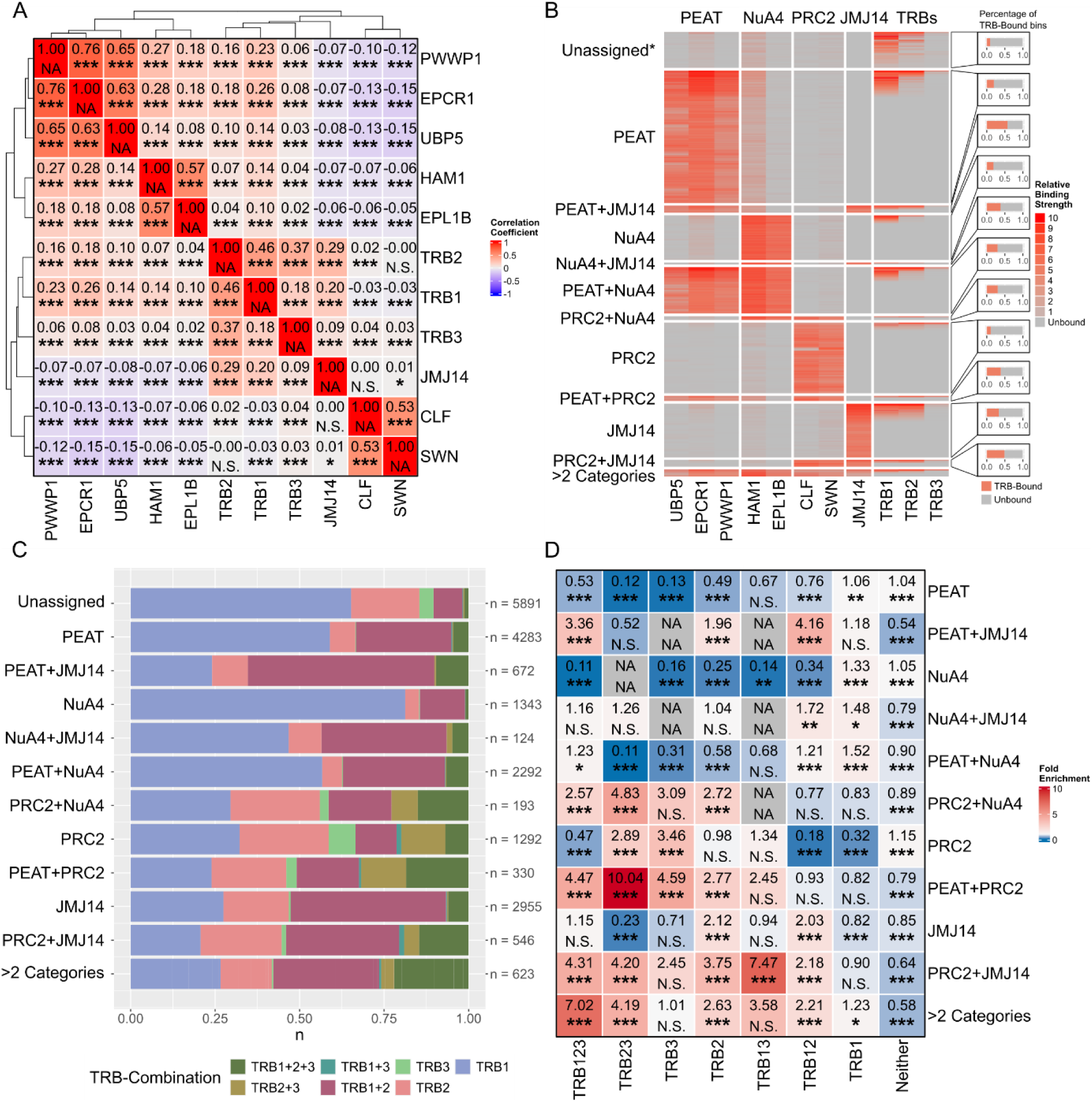
Analysis of genomic binding locations of eleven epigenetic regulatory proteins. **A)**; Pairwise Pearson correlation matrix of ChIP-Seq peaks derived from eleven regulatory proteins and assigned to 500 bp genomic bins. Significance levels: ***, p<=0.0005, **, p<=0.005, p<=0.05, N.S., p>0.05. **B)**; Left, Heatmap depicting the genomic bins bound by each of the ChIP-Seq sets used in A. Relative binding strength of each peak expressed through deciles. Columns depict proteins grouped by regulatory complex and rows depict bins assigned to complexes based on presence of at least two complex components. Bins assigned to more than two complexes were grouped into one category. *, 58676 bins which were neither assigned to complexes nor TRB-bound were excluded for the sake of readability. Right, Percentage of TRB-bound bins for each category. **C)** Distribution of TRBs in the TRB-bound bins of each category assigned in B. **D)** Pairwise exact test statistics as described by Wang et al. (2015) for bins assigned to complex categories and their corresponding TRB combinations. Significance levels: ***, p<=0.0001, **, p<=0.001, p<=0.01, N.S., p>0.01.

A heatmap using the matrix of decile transformed binding sites across target categories underscored that TRBs rarely bind chromatin without any of the described partner complexes (Figure 2B). The heatmap visualized the mostly exclusive, non-overlapping nature of complex categories except for PEAT and NuA4, which are equally likely to bind chromatin in combination or alone. Next, we evaluated if TRB paralogs showed preference for specific partner complexes. Association of fully resolved TRB binding categories indicated a high proportion of PEAT and NuA4 sites that were exclusively bound by TRB1 (Figure 2C). If JMJ14 was also associated with PEAT and/or NuA4, the proportion shifted to include more peaks co-bound by TRB1 and 2 or by all TRBs. The shift in distribution was significantly different from the expected based on a SuperExact Test of expected combinations (Figure 2D) [25]. Presence of TRB2, in particular in combination with TRB1, was most significantly enriched for JMJ14 containing bins, irrespective of the presence or absence of other interacting complexes. In contrast, PRC2 was most overrepresented at peaks only bound by TRB3, followed by the combination of TRB3 and TRB2. At the rare sites where PRC2 co-bound with JMJ14 or with PEAT, the overrepresentation of TRB1/TRB3 and TRB2/TRB3 peaks became much higher.

Since TRB1 and TRB3 showed a strong propensity to associate respectively with PEAT/NuA4 and PRC2, we performed Immunoprecipitation Mass Spectrometry (IP-MS) to evaluate whether preferred complex associations could also be observed at the protein level. Plants expressing *TRB1* or *TRB3* to *YFP* fusions under the control of the *Cauliflower Mosaic Virus 35S* promoter (*35Sp*) were immunoprecipitated from nuclear extracts using a GFP-trap resin. Samples prepared from *35Sp-ENHANCED DISEASE SUSCEPTIBILITY 1 (EDS1)-YFP* transgenic lines were used as background control. Using biological triplicates, overall 1149 proteins were identified, of which 96 were significantly enriched in either TRB1 or TRB3 pull-downs or both (Supplemental File 4). Co-purified proteins included components of PEAT, NuA4, JMJ14 complexes and PRC2. Only 6 and 7 proteins were significantly enriched in TRB1 over TRB3 and TRB3 over TRB3, respectively (Figure 3A). PEAT-components EPCR1 and EPCR2 showed significant preference to purify with TRB1 over TRB3 while other PEAT components showed a trend towards a higher enrichment for TRB1 (Figure 3B). In contrast, the only NuA4 component that was enriched, TRA1A, was equally enriched by both TRBs (Figure 3C). Equal purification was also observed for JMJ14 components, which showed only a slight bias towards TRB3 (Figure 3D). EMF1 was the only PRC2 associated protein identified and it was exclusively enriched by TRB3 (Figure 3E). Both, TRB1 and TRB3 co-enriched the transcription factor (TF) VIVIPAROUS1/ABI3-LIKE1;VP1/ABI3-LIKE 1 (VAL1), which can recruit PRC1 and PRC2 to target regions.

**Figure 3.**
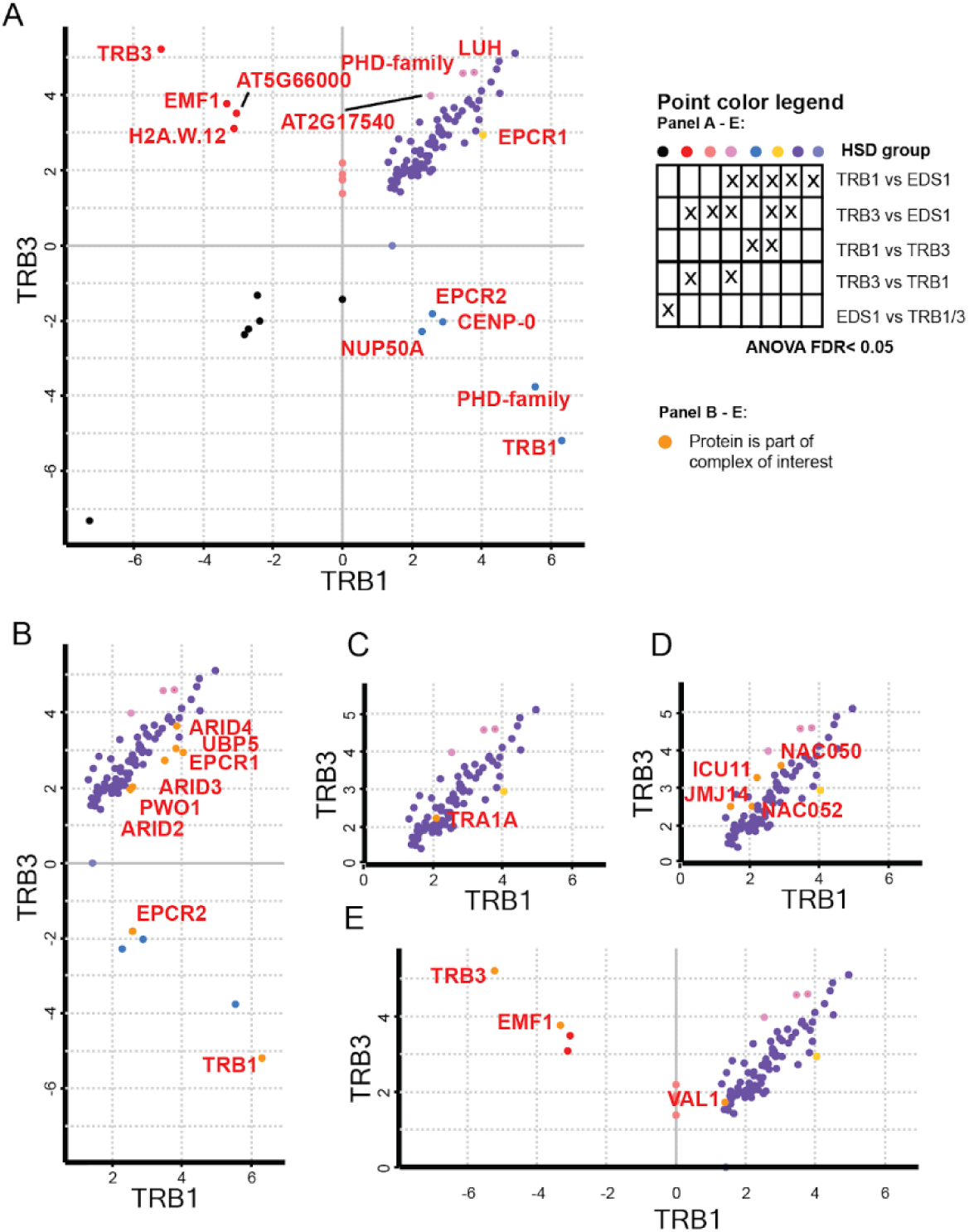
TRB1 and TRB3 interactome. **A)** Comparisons of IP-MS between TRB1 and TRB3 co-purifying proteins in nuclear extracts of 14-day-old seedlings. Axes show log2-fold enrichment data of log transformed LFQ data from TRB1-GFP vs EDS1-GFP and TRB3-GFP vs EDS1-GFP as indicated). Values with statistical significance (FDR < 0.05) enrichment or depletion as determined by ANOVA and HSD test are indicated. Significance groups indicated by colors as indicated in the legend. Gene symbols are indicated for genes that show significant difference between TRB3 and TRB1. **B)** Excerpt of A) showing PEAT complex components as indicated by color and gene symbol. **C)** excerpt of A) for NuA4 complex. **D)** excerpt of A) for JMJ14 complex. **E)** as in A for PRC complex.

### TRB1-3 peaks enrich distinct DNA motifs dependent on co-bound regulatory complexes

Analysis of DNA motif enrichment at TRB1-3 peaks classified by co-associated complexes provided further evidence of specialization among the three TRB paralogs. We used the XStreme pipeline of the MEME-Suite to evaluate DNA motif enrichment at TRB1, 2 and 3 peaks as well as the peaks assigned to the different complex categories for each paralog (Figure 4, Supplemental File 5). A comparison of enriched motifs of all TRB1-3 peaks regardless of complex association revealed that the main telo-box motifs bound by TRB1-3 are not identical. While all three paralogs enrich the canonical “AAACCCT” telo-box motif, TRB1 also significantly enriched the similar but distinct “CRACCTA” motif. While multiple secondary motifs were found for all three paralogs, it stands out that motifs associated with the basic/helix-loop-helix (bHLH) transcription factor family are only enriched in TRB1 ChIP-Seq peaks. Motifs related to TCP-TFs, on the other hand, only enriched in the peaks of TRB1 and TRB2.

**Figure 4.**
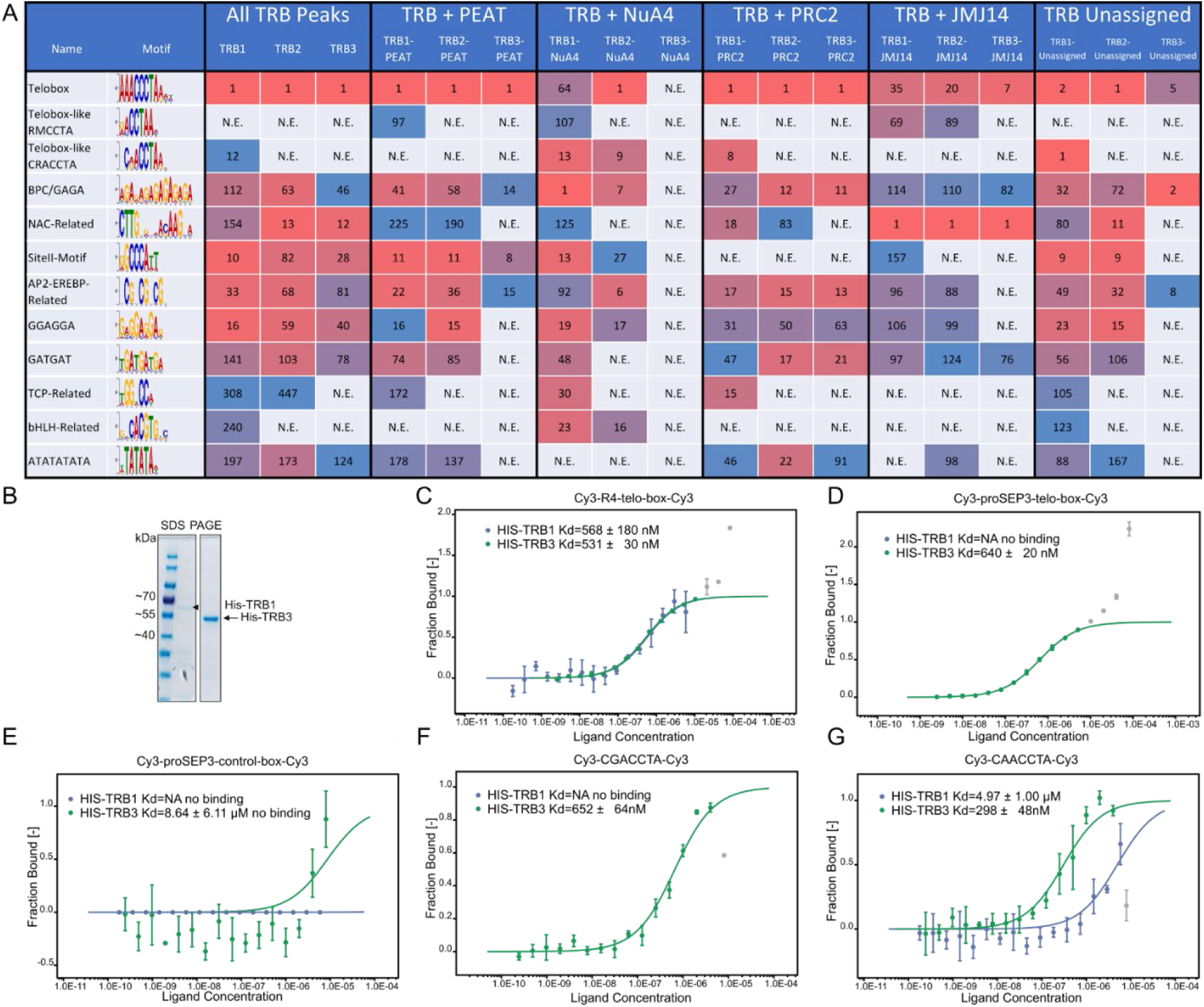
Analysis of TRB bound and associated motifs. **A)** DNA motifs enriched in ChIP-Seq peaks of TRB1, TRB2, and TRB3. Either all peaks, peaks assigned to gene regulatory complex binding sites or only peaks without complex assignment were used for the analysis as indicated in the header. Numbers indicate enrichment rank in the MEME-Suite results. **B)** Bacterial expressed HIS-TRB1 and HIS-TRB3 after affinity purification and SDS-PAGE. **C)** Fraction of Cy3-labelled double stranded 4x repeated telo-box probe bound by HIS-TRB1 and HIS-TRB3. **D)** as C) using Cy3 lablelled telo-box motif as found in the promoter of SEP3. **E)** as C) using a Cy3-labelled control region from the SEP3 promoter **F)** and **G)** as C using Cy3 labelled double stranded oligos containing the CGACCTA and CAACCTA motif, respectively.

To identify motifs associating TRBs with one of the four regulatory complexes, we repeated the XStreme analysis pipeline using only TRB peaks from sites that were categorized as exclusive binding locations of PEAT, NuA4, PRC2, or JMJ14. This detailed analysis showed that TRB1 enriches a second telo-box-like motif (“RMCCTA”) at sites not co-bound by PRC2. It furthermore indicates that the BPC-transcription-factor-related GAGA-box and NAC-TF associated motifs are universally enriched at all sites, independent of co-bound complex components. Other secondary motifs are specifically enriched in sites co-bound by certain regulatory complexes. This includes the siteII motif, which does not enrich at sites with PRC2 binding, ATATAT, which does not enrich at sites assigned to NuA4, and the contrasting bHLH-related motifs, which exclusively enrich at sites with NuA4 binding. Looking closer at the ranks associated with each enriched motif reveals interesting differences between sites with otherwise similar enrichment patterns. Most strikingly, peaks associated with sites of all three paralogs co-bound by JMJ14 show NAC-related TFs as their highest ranking motif, even surpassing the ranks of telo-box motifs.

These results indicate a substantial difference in DNA motifs found at sites where TRBs bind together with PEAT, NuA4, PRC2, or JMJ14 and could therefore indicate the role of different TFs in determining the targeting of epigenetic regulatory complexes.

### TRB1 binding to non-canonical telo-box motifs is not explained by affinity to single motifs

Since differences in the telo-box related consensus motifs suggested that the DNA binding properties may differ between TRB1 and TRB2/3, we tested the affinity of TRB1 and TRB3 to single telobox and the two variants of the “CRACCTA” motifs by microscale thermophoresis (MST). MST determines dissociation constants between fluorescently labelled targets and unlabelled ligands by measuring changes in the velocity by which the fluorophore moves in or out of 1-6° K temperature gradients (Seidel et al. 2013).[26]. TRB1 and TRB3 were previously shown to bind probes containing four teloboxes (R4) that imitate telomeric repeats *in vitro* [27].

We could reproduce binding to R4 probes for bacterially expressed six histidine (HIS)-tagged TRB1 and TRB3 with binding affinities that were in the range of those previously reported (K_D_=567± 180nM and K_D_=531± 30nM for TRB1 and TRB3, respectively) (Figure 4B and C). In contrast, only HIS-TRB3 interacted with an 28bp oligomer derived from the *SEP3* promoter containing a single telobox (K_D_= 640 ± 19 nM) (Figure 4D and E). Since the affinity of HIS-TRB3 towards the single telo-box motif was comparable to that of the R4 telobox repeat, a non-cooperative interaction mechanism between TRB3 and DNA is probable. Motif enrichment suggested that TRB1 may bind to single CRACCTA motifs; however, HIS-TRB1 did not bind to the CGACCTA variant and only with low affinity to the CAACCTA variant (K_D_ = 4.9 ± 1.0 µM) (Figure 4G and G). Unexpectedly, HIS-TRB3 bound CGACCTA with similar affinity as single teloboxes (CGACCTAA: K_D_ = 652 ± 64 nM) and CAACCTA slightly better (K_D_ = 298 ± 48 nM) (Fig. 4F and G).

In sum, TRB1 seems unable to bind single telo-box and CRAACTA motifs *in-vitro,* indicating that *in vivo* enrichment of these motifs by TRB1 may depend on the presence of accessory factors.

### Genes bound by different regulatory complexes participate in distinct biological processes, dependent on co-bound TRBs

Next, we investigated whether the epigenetic regulatory complexes diverged not only in their binding sites and TRB paralog association, but whether their target genes were also involved in different biological processes. Through the annotatePeak function of the ChIPseeker library, peaks were annotated to the *A. thaliana* genome. This revealed that, while only 8.6% of the 500-bp bins were bound by TRBs, 29.4% of all annotated nuclear genes exhibited binding of at least one TRB.

We started by identifying the most enriched GO-terms for each complex, regardless of the TRB paralog they were associated with (Figure 5 A and Supplemental File 6). Overall, it was possible to assign specific functions to each complex combination. Ribosome-related GO-terms were significantly enriched in genes assigned to PEAT with and without NuA4; however, the significance was higher for PEAT only genes, indicating a previously undescribed role of the PEAT complex in ribosomal regulation. PRC2 assigned genes enriched terms related to floral development. This is consistent with previously published functions of TRB-PRC2 complexes [14]. NuA4-assigned genes with and without PEAT overrepresented terms annotated to the circadian clock, an association that was previously detected for NuA4 [28]. The combination of JMJ14 and NuA4 target genes enriched GO-terms related to immune responses. Last, the five GO-terms with most enrichment in the unassigned set of genes were all related to oxygen response/hypoxia, indicating that our current understanding of TRB-related gene regulation is still incomplete.

**Figure 5.**
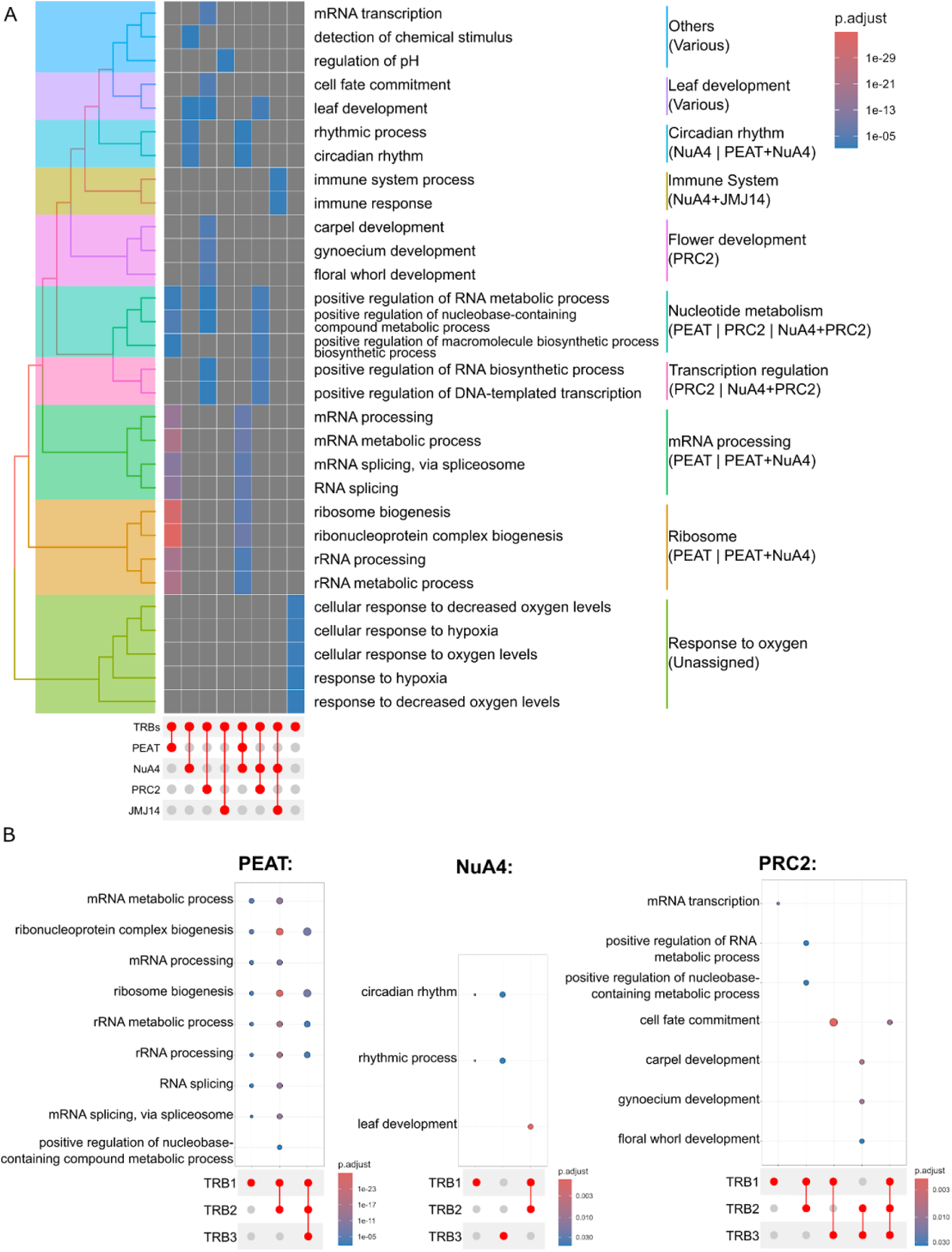
Gene Ontology enrichment analysis. **A)**, Enriched GO-Terms for genes assigned to either no complex (Unassigned), PEAT, NuA4, PRC2, JMJ14, or any of the four combinations of two complexes. Only the 5 most enriched GO-Terms of the “Biological Process” aspect are shown. Similar terms are summarized on the right. **B)** Enriched GO-Terms for the gene sets assigned to PEAT, NuA4, or PRC2, separated by co-bound TRB-paralog. The same terms as in A are shown.

To expand this analysis and differentiate complexes based on their co-bound TRB paralog, we determined GO-term enrichment of all combinations. Overall, the GO-terms barely correlated between genes assigned to the same complex, but bound by different TRB homolog combinations (Supplemental Figure 4). This functional split constitutes further evidence of the specialization between TRB paralogs, even within the same complex.

Since epigenetic regulatory complexes enriched different GO-terms, based on which TRB paralog combination they associated with, we took a closer look at the GO-terms enriched for genes assigned to PEAT, NuA4, and PRC2 (Figure 5B). Comparing the TRB-paralog combinations, it became clear that the affinity of PEAT towards ribosomal and RNA-processing genes is driven largely by TRB1, as almost all possible TRB1-containing combinations enrich these terms. The role of NuA4 in the circadian clock appears to be coinciding with exclusively TRB1 or TRB3 bound genes. PRC2 target genes involved in floral development are only enriched in the genes that are also bound by TRB2 and TRB3.

### Transcriptomic changes in *trb* mutants are partially explained by the assigned regulatory complexes

In order to validate the complex assignments and to improve our understanding of the effects of TRBs on gene expression, we tested whether specific complexes are over- or underrepresented in the sets of genes assigned to PAM-clusters identified for DEGs in single and double *trb* mutants (Figure 1E). The normalized average expression profile across all genes per cluster confirmed that clusters 1 and 2 exhibit an inverse pattern, with the loss of *TRB1* appearing as the driving factor behind the effects (Figure 6A). Genes in cluster 4 showed a progressive increase of expression, with mutations in *TRB2* and *TRB3* as strongest drivers. Based on these patterns, we hypothesized that cluster 1 contained genes primarily controlled by the activating PEAT complex, while cluster 3 genes were likely controlled by PRC2. Using the gene-to-complex assignments produced for the gene ontology analysis (Supplemental data 3), we intersected the PAM-clusters with complex combinations and performed multi-set interaction analysis according to [25] (Figure 6B). The test confirmed that cluster 1 exhibits a 2.2-fold enrichment of TRB-bound genes assigned to PEAT. Genes assigned to both PEAT and JMJ14, however, were significantly depleted in the same set (0.2-fold). Interestingly, cluster 2 exhibited an inverse enrichment, in line with its inverse expression profile. In this cluster, all combinations that include JMJ14 but not PRC2 were significantly enriched (between 1.8 and 3.2-fold enrichment). This suggests a so far undescribed role of PEAT and/or NuA4 in repressive gene regulation when found alongside JMJ14. Cluster 3 also shows a 2.9-fold enrichment of genes assigned to only JMJ14, but without a clear indication on how the expression profile can be explained. Lastly, cluster 4 is characterized by a 2.8-fold enrichment of PRC2-assigned genes with a simultaneous 0.3-fold depletion of PEAT-assigned genes, as expected based on its expression profile.

**Figure 6.**
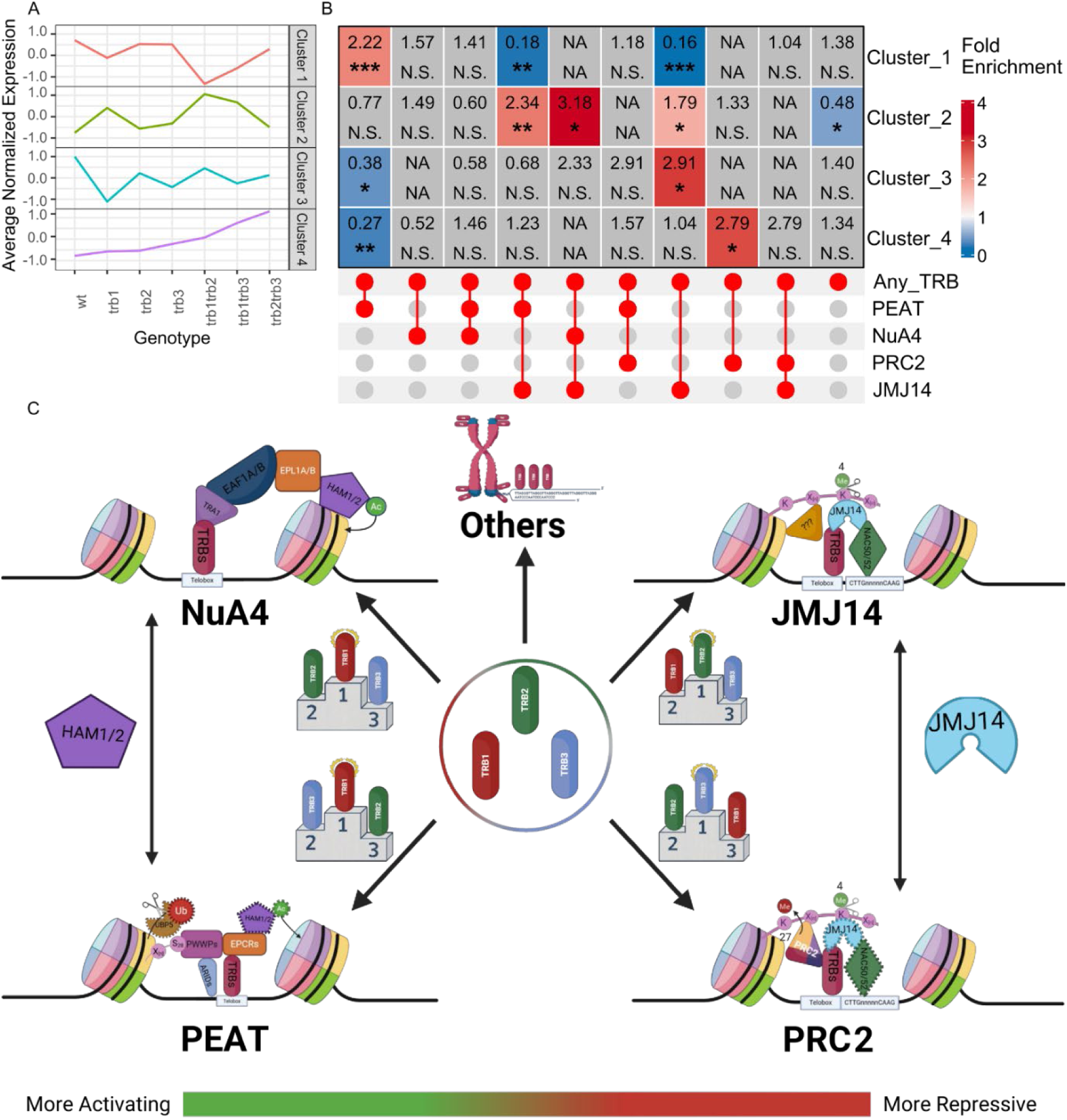
TRB-mediated gene regulation in epigenetic context. **A)** Average normalized expression of the transcription clusters displayed in Fig. 1E. Values of three replicates of each of genotypes were averaged. **B)** Pairwise Exact test statistics as described by Wang et al. (2015) for the same clusters as A intersected with the sets of genes annotated to either one, two, or none of the epigenetic regulatory complexes. Significance levels: ***, p<=0.0005, **, p<=0.005, p<=0.05, N.S., p>0.01. **C)** Model of TRB-mediated epigenetic gene regulation. Complex components which are not core to the TRB□containing complexes are indicated with dashed outlines. TRB-paralog specialization is conveyed via podiums. “???” indicates potential unknown complex participants. HAM1/2 and JMJ14 are shared between the complexes on the activating and repressive side, respectively.

## Discussion

### TRBs associate with distinct epigenetic regulatory complexes in a primarily mutually exclusive manner

Since it was first discovered that TRBs can participate in multiple epigenetic regulatory complexes, it had been unclear whether TRB-bound genomic sites act as “landing sites” for multiple complexes, or whether each site is associated with only a single complex. Our analysis of the ChIP-Seq-determined binding sites of eleven complex-forming proteins (including two novel TRB sets) serves as a useful tool to evaluate the overall binding behavior of TRBs. According to our analysis, it is likely that the majority of TRB-binding sites are associated with a single regulatory complex (Figure 2). The only exception seems to be PEAT and NuA4, which share a significant proportion of their binding sites, likely caused by the role of HAM1 in both complexes [12, 29]. It should, however, be noted that all the ChIP-Seq datasets included in this study are obtained from whole seedlings. It is therefore likely that temporal and/or tissue-specific binding differences are obscured in our analysis. In addition to mutual exclusivity, we also discovered that a large portion of the binding sites of the four complexes included in this study were not bound by any of the three TRB paralogs. In the case of PEAT, this is unexpected, since TRBs have been proposed as core components [12]. In part, this finding may be explained by differences in stringency thresholds defining binding sites in different bioinformatics pipelines.

Our analysis provides strong evidence of additional, thus far undescribed, TRB-containing complexes. First, the presence of a large contingency of TRB-JMJ14 co-binding sites in the absence of other complex components strongly indicates the presence of a JMJ14-mediated histone modification complex that acts independently of PRC2, while maintaining JMJ14 association with TRBs. NAC50/52, two strong TRB interactors with slight preference for TRB3 over TRB1, have already been shown to form regulatory complexes with JMJ14 [22]. Furthermore, the evidence clearly shows a split between PEAT and NuA4 bound TRB-sites, expanding on the previously observed role of TRBs in PEAT complexes which include the HAM1 component of NuA4 [11]. This serves as evidence of the presence of an additional NuA4-TRB complex targeting sites independent of PEAT. In addition to their genomic binding, evidence of these two proposed complexes can also be seen in the IP-MS data. TRA1, which had previously been described as the TF-binding domain of the NuA4 complex [29], is the NuA4 complex component with the strongest TRB1/3 binding score (Figure 3). Last, a substantial number of target sites is associated with TRBs in absence of all other chromatin complex representatives. It is to be expected that other TRB partners will be identified to co-occur at these sites.

### The three paralogs TRB1, 2 and 3 specialize in specific roles, without losing their redundancy

Although reverse genetics suggests almost complete redundancy between the three paralogs TRB1, 2, and 3, whether this redundancy is replicated on a molecular level had never been studied in detail. On the molecular level, TRBs exhibited an unexpectedly strong degree of specialization. In particular, a clear split between TRB1 and TRB2/3 could be detected throughout the analysis. First, transcriptome changes of single and double *trb* mutant combination identified two sets of genes with opposing expression patterns that were predominantly associated with the loss of *trb1* (Figure 1 and Figure 6). A different set of genes was higher expressed when the *trb3* alles was combined with *trb2* or *trb1*. Furthermore, the association between TRB1 and activating complexes such as PEAT and NuA4 could be seen in the overrepresentation of TRB1 bound sites in the ChIP-Seq data of these complexes; furthermore, it is corroborated by GO-term analysis since terms enriched for TRB1 a more significantly overrepresented for TRB-PEAT and TRB-NuA4 target genes (Figure 2 and 5). Conversely, the same data indicate that TRB2/3 associate more strongly with the repressive complex PRC2 and the proposed JMJ14-containing complex. The same trend can be observed in the IP-MS data comparing TRB1 and TRB3 co-purifying proteins since the only identified PRC2-associated protein, EMF1, was exclusive to TRB3 while PEAT-component EPCR1 was only co-purified with TRB1 (Figure 3).

### TRBs are low affinity transcription factors explaining their dependence on other interaction partners and motifs for target association

Although TRB proteins can bind telo-box and related motifs, their affinity towards tandemly repeated motifs was moderate at best and weak to non-binding in the case of single telo-box or related motifs (Figure 4). Hypothetically, high affinity binding to telo-box motifs would strongly tether TRBs to telomeres, possibly preventing their binding to single motifs at genic regions. On the other hand, the weak affinity of TRBs to target sites *in vitro* suggests that TRBs require assistance to associate with genic target regions *in vivo*. TRBs feature a H1/H5-related domain that may increase the affinity to well-positioned telo-box motifs by interacting with core nucleosomes. Previous studies have shown that presence of the linker histone H1 competes with TRB1 binding to telomere-derived regions [8]. Alternatively, the association with chromatin complexes and other TFs may stabilize TRBs at single telo-box motifs.

While the analysis of DNA motifs could not fully explain likely (co)-recruiting motifs for different TRB-associated complexes, it provided further evidence of specialization. A few DNA motifs show differential enrichment between the regulatory complexes (e.g., the site II motif and bHLH related motifs) and ranks of the enriched motifs differ substantially between the complex target sites (Figure 4). Most obviously, sites of TRB-JMJ14 binding enrich NAC-related motifs to a stronger degree than telo-box motifs. Several studies had already presented interaction data identifying a complex of NAC50/52 with TRBs and JMJ14 [22, 23], which was also confirmed in our study (Figure 3). The predominant enrichment of NAC motifs indicates a TRB recruitment mechanism as part of the TRB-NAC-JMJ14 complex involving NAC’s affinity to their cognate motifs.

In a second case, interaction between two closely linked motifs appears not to rely on a common complex. Earlier studies revealed that telo-box motifs in combination with site II motifs are able to act as transcriptional enhancers at ribosomal protein encoding genes [30]. Our analysis showed that TRB-PEAT bound genes enrich site II motifs and are associated with GO-terms associated with ribosomal functions (Figure 2 and 4). In contrast, the site II motif binding TEOSINTE BRANCHED1/CYCLOIDEA/PROLIFERATING CELL FACTOR (TCP) family TFs were not enriched in our IP-MS data (Figure 3).

Last, no singular motif was found that only enriches at sites co-occupied by PRC2 and TRBs. While GAGA-motifs, which are bound by TFs of the BPC family, were previously proposed a co-recruiters of PRC1 or PRC2 components [19, 31, 32], the corresponding motifs are also enriched at TRB-PEAT and TRB-NuA4 leaving the question of how PRC2-TRB sites are defined unanswered for now. BPC TFs were not enriched in our IP-MS dataset.

In summary, we observed an overall trend of TRB specialization, which can be best described as a preference for specific sites and complex partners (Figure 6C). TRB1 appears to have a strong affinity with PEAT and NuA4, but is also present at many unrelated binding sites. By contrast, TRB3 and TRB2 are predominantly, though not exclusively, found at PRC2 and JMJ14 locations, respectively. These results are consistent with the remarkable redundancy observed at the phenotypic level. It is possible that TRBs are at the beginning of a process of functional specification. Alternatively, their partial functionalization could provide robustness to epigenetic gene regulation, particularly if changes in their relative abundance and activity are connected to environmental or developmental cues. Further studies are required to answer these questions.

## Materials & Methods

### Plant material and growth conditions

Plants were grown in greenhouse conditions or growth chambers as indicated under long day (LD) (16h light, 8h dark) photoperiod at 22°C ambient temperature. Plants were randomized within trays for phenotypic analyses. Plants for RNA-seq analysis were grown in growth chambers. Three biological replicates were grown in 1 week intervals in the same chamber, material was collected at ZT10 from 14-day-old seedlings. For ChIP-seq, plants were grown in tissue culture on GM plates in LD conditions at 21°C. Material from replicated plates was collected at ZT10 as biological replicates. The *trb1-2* (Salk_001540) and *trb3-2* (Salk_134641) alleles are previously described T-DNA insertion lines, *trb2-2* and *trb2-3* are CRISPR/Cas9 edited alleles as previously described except that the editing transgene was removed by segregation [14]. Double and triple mutants were generated by crosses. Transgenic lines TRB2pro-TRB2-YFP and TRB3pro-TRB3-YFP were previously described [14].

### Scoring of flowering time

Flowering time was scored as the number of leaves at the main shoot (rosette and cauline leaves). Statistical analysis was done by ANOVA with HSD grouping using the agricolae package in R. To distinguish segregating *prope* triple from double mutants, genotyping of individual plants was carried out on genomic DNA prepared using Biospring96 (Qiagen) using manufacturer’s instructions. Alleles trb1-2 and trb3-2 were amplified using SALK_LBb1.3: ATTTTGCCGATTTCGGAAC in combination with SALK_001540_RP: ATGCCACCACAATAAATCTCG and SALK_13464_RP: ATGGTTCACGAGAAACCTGTG, respectively. To distinguish between trb2-3 and TRB2, two reactions were carried out for 28 cycles at 62°C annealing temperature dCAPS_TRB2_R: ATTGCCTCAAAGATGATCTTATCC in combination with 8-18-10-specifi: ACTTCCCCCGGAGGTTCTTG and 8-18-10-WT: ACTTCCCCCGGAGGTTCTG, respectively.

### RNA preparation and RNA-seq

Total RNA was extracted from 3-4 14-d-old-seedlings with an RNeasy® Plant Mini kit (Qiagen) according to the manufacturer’s instructions. To remove gDNA contamination, 10 µg of total RNA was DNase I treated, using the DNA-freeTM DNA Removal kit (InvitrogenTM), as described in the kit’s instructions. RNA quality was assessed by Agarose Gel electrophoreses of an 200ng aliquot DNaseI treated RNA. The RNA samples were sent to BGI TECH SOLUTIONS (HONGKONG) for poly-A enrichment, library preparation and directional paired-end Nanoball sequencing on the DNBSEQ platform.

### RNA-seq analysis

Paired end reads were mapped to the Arabidopsis thaliana TAIR10 reference genome indexed with the Araport11 genome annotation using STAR. Read counts were pooled for all splice variants as per gene counts. Sense strand gene counts were used for differential expression analysis with the R package DESeq2 using a threshold of padj < 0.05 to set differential expression of mutants vs Col-0. Venn diagrams and statistical testing of overlaps between samples used R packages ggvenn and SuperExactTest, respectively. Clustering of expression data and drawing of gene-normalized expression heatmaps were carried out using R package ComplexHeatmap using PAM-clustering.

### ChIP and ChIP-seq library preparation

For all ChIP experiments, 2 g of 14-d-old seedlings were collected in 50 ml 1x PBS buffer (137 mM NaCl, 1.8 mM KH_2_PO_4_, 10.1 mM Na_2_HPO_4_, 2.7 mM KCl), fixed with 1 % formaldehyde under vacuum two times for 10 min after which the crosslinking reaction was quenched with 5 ml glycine (1 M) under vacuum for 5 min. Fixed plant material was collected in a sieve, washed with autoclaved water, and dried with paper towels before being snap frozen with liquid N_2_. Frozen samples were ground at 7200 rpm three times for 30 s, using the Precellys Evolution Homogenizer in combination with a Cryolys Cooling Option (Bertin Instruments) in 7 ml reaction tubes with 3mm ceramic beads.

To extract nuclei, the ground samples were mixed with 30 ml NIB buffer (50 mM HEPES-NaOH (pH 7.4), 5 mM MgCl_2_, 25 mM NaCl, 5 % sucrose, 30 % glycerine, 0.25 % Triton X 100, freshly add: 0.1 % β-mercaptoethanol, 0.1 % SIGMA proteinase inhibitor), vortexed, filtrated using Miracloth (Merk) and spun down at 4000 rpm and 4 °C for 10 min. The pellet was resuspended in 20 ml 1x Washing buffer (16.7 mM HEPES-NaOH (pH 7.4), 6.7 mM MgCl_2_, 33.3 mM NaCl, 13.3 % sucrose, 13.3 % glycerine, 0.25 % Triton X 100, freshly add: 0.001 % β-mercaptoethanol, 0.001 % SIGMA proteinase inhibitor) and spun down at 4000 rpm and 4 °C for 10 min. Then, extracted nuclei were resuspended in TE-SDS (1 mM EDTA (pH 8.0), 10 mM Tris-HCl (pH 7.4), 0.25 % SDS) in a total volume of 600 µl, rotated at 4 °C and 12 rpm for 20 min, split in 2x 300 µl and sonicated with a Bioruptor Sonicator (Diagenode) that was attached to a Minichiller cooling system (huber) (Programme: red - 0.5 (on); green - 1 (off); 15 min, H) to produce DNA fragments of ∼200-500 bp. Sonicated chromatin was separated from debris by centrifugation at 4 °C and maximum speed for 10 min. For ChIP-seq, 400 µl sonicated chromatin were mixed with 600 µl of IP dilution buffer (80 mM Tris-HCl (pH 7.4), 230 mM NaCl, 1.7 % NP40, 0.17 % DOC), 2 µl RNase I (10 mg/ml), 2 µl DTT (1M), and 2 µl SIGMA proteinase inhibitor. Afterwards, equal volumes of the sonicated chromatin mix were split into two different tubes, 5 µl of α-GFP (ab290, Abcam) were added to carry out IP. Samples were rotated at 4 °C, 12 rpm overnight in a bohemian wheel. After overnight incubation, unspecific precipitates were removed by centrifugation (4 °C, 20000g, 10 min) and the supernatant transferred to a tube containing 30 µl rProtein A Sepharose Fast Flow antibody purification resin (GE Healthcare) beads equilibrated in RIPA buffer (0.6x IP Dilution buffer, 0.1 % SDS). Samples were rotated at 12 rpm and 4 °C for 3 h. After centrifugation, 200 µl of the supernatant from control samples was reserved as input and kept on ice. Beads were washed with 1 ml RIPA for five times to remove the background. At the 5^th^ time, the samples were transferred to fresh tubes with 800 µl RIPA and protein-DNA complexes were eluted from precipitated beads by mixing them two times with 160 µl glycine elution buffer at RT. IP samples were neutralised with 80 µl of Tris-HCl (1 M, pH 9.7). IP samples were de-crosslinked by adding 8 µl SDS (10 %) and 5 µl proteinase K (5 mg/ml). For input samples, only 5 µl proteinase K was added. DNA was extracted twice with equal amounts of phenol/chloroform and precipitated with 1/10 volumes NaAC (3 M), 2.5 volumes EtOH (100 %), and 1 µl glycogen (10 mg/ml) at −20 ° for 3 h. Afterwards, the DNA was washed with 1 ml EtOH (70 %), dried, and resuspended in 14 µl H_2_O.

For ChIP–seq library preparation, two independent immunoprecipitations for Col-0, TRB2pro-TRB2-YFP and TRB3pro-TRB3-YFP were processed. Libraries were prepared with Ovation Ultralow Library System (NuGEN) according to the manufacturer’s instructions, using 71% (10 µl) of each ChIP as starting material. Before amplification DNA concentration was measured, using a Qbit 4 (ThermoFisher Scientific), to determine the appropriate number of PCR cycles needed for each sample (see manufacturer’s manual). After amplification, DNA was run on a 2 % low-melt agarose gel and fragments between 200 and 500□bp length were purified using the MinElute Gel Extraction Kit (Qiagen) according to the manufacturer’s instructions except that gel fragments were solved at RT and eluted in 15 µl EB buffer. An aliquot of each library was tested via qPCR before and after PCR amplification to confirm that libraries showed similar fold-change between control and target regions. Sequencing was performed as single-end 100-nt reads (ca 13 mio reads/sample) on the Illumina HiSeq3000 platform by the Max Planck Genome Centre Cologne.

### ChIP-seq analysis

After sequencing, adapter sequences ≥ 12 bp were removed using Cutadapt [33]. Reads were aligned to the *A. thaliana* genome (TAIR10) with the Burrow-Wheeler Aligner (BWA) [34] and BAM-files created using SAMtools [35]. SAMtools was used to remove multi-mapping reads by filtering with MAPQ score < 10, which resulted in 8.8 to 12.2 million reads per sample. Unique BAM-files were indexed with SAMtools, normalised to Counts Per Million mapped reads (CPM), and converted to bigWig-files using bamCoverage of the deeptools2 suite [36] for visualization in the Integrated Genome Viewer (IGV). A blacklist of over- and under sampled regions was generated by scoring read coverage of input and Col-0 ChIP samples across 200bp windows using BEDtools [37]. Windows that were statistical outliers were determined using R and subsequently removed from the analysis. EPIC2 was used to determine enriched regions in two replicates against the pool of two Col-0 control IPs using pooled input samples as correction [38]. Replicates were compared using the Irreproducible Discovery Rate (IDR) framework [39]. Peak passing the threshold of 0.01 > IDR were merged using bedtools. Previously generated 35Sp-TRB1-YFP reads were included in the IDR analysis for better comparison [9].

### Analysis of the binding behavior of various epigenetic regulatory complexes

Binding peaks of UBP5, EPRC1, PWWP1, HAM1, and EPL1B were sourced from [11], CLF and SWN from [21], and JMJ14 were obtained from [23]. To visualize the overlapping binding sites, the TAIR10 genome of *A. thaliana* was tiled into 238296 bins of 500 bp length and all peaks of these eight datasets and TRB1,TRB2 and TRB3 were assigned to overlapping bins. As the datasets were derived using different ChIP-Seq pipelines, the “Score” column of each dataset was normalised into deciles. Preliminary analysis of the overlap of the ChIP-Seq sets was performed through pearson correlation using Hmisc [40].

To visualize the overlapping binding sites, the bins were assigned to distinct categories: Bins bound by at least two of the PEAT components, both NuA4 components, both PRC2 components, or JMJ14 were assigned to “PEAT”, “NuA4”, “PRC2”, or “JMJ14” respectively. In addition, each of the 12 possible combinations of multiple complex assignments were added along with a category for unassigned bins, bringing the total to 17 categories. Each bin was assigned to one of these categories. Statistical analysis was performed using the MSET function of the SuperExactTest package [25] for pairwise comparison of the overlap of the generated categories with TRB1,2,3-bound bins. Heatmaps were generated using the ComplexHeatmap [41] package.

### Cis-motif enrichment analysis

DNA motifs enriched in peak-assigned bins were identified through the XSTREME pipeline of the MEME-suit [42] using standard settings except for *--meme-mod “anr”*, providing the DNA motifs identified by [43]. Motifs were declared as telobox-like, if the identified motif was closely related to the telobox, but did not fully capture the canonical *Arabidopsis* telomere repeat sequence of TTTAGGG. For each peak category, motifs were ranked based on their e-Value.

### Gene Ontology enrichment analysis

The peaks of all ChIP-Seq sets used in this study were annotated to genes using “annotatePeak” of the “ChIPseeker” package [44, 45]. Since epigenetic regulatory complexes are not solely found at or near the TSS, the parameters “tssRegion” and “overlap” were set to “c(−2000, 2000)” and "all" respectively in order to correctly assign binding sites further away from the TSS. The resulting genes were assigned to epigenetic regulatory complexes in the same manner as the genomic bins. The different sets were subsequently used to calculate GO-Term enrichment using the “clusterProfiler” package [46].

### Sample preparation for LC-MS/MS

Leaf tissue (7g) harvested from 5-week-old transgenic plants (*CaMV 35Sp-TRB1-GFP*, *CaMV 35Sp-TRB3-GFP*, *CaMV 35Sp-EDS1-GFP*) was cut with scissors into 0.5 - 1.0 cm pieces and disrupted on ice in 15 ml Precellys® tubes containing 5 ml extraction buffer (2 M hexylene glycol, 0.5 M PIPES-KOH pH7.0, 10 mM MgCl2, 5 mM beta-mercaptoethanol) and 13-15 sterilized metal beads using a Precellys® 24 homogenizer (Bertin instruments) for three rounds set to 10 s at 7500 rpm. Samples were filtered through a single and then a double Miracloth (Merk) layer, adjusted to a volume of 45 ml. 10% Triton X-100 was added stepwise to a final concentration of 0.8%. While samples were incubated on ice, the Percoll® (Sigma-Aldrich) gradient was assembled by carefully underlying 6 ml of 30% Percoll® solution with 6 ml of 80% Percoll® in a centrifuge tube (Beckman Coulter #355631). In parallel, three 15 ml aliquots per sample were layered onto gradients and centrifuged (2,000 g, 4 °C, 30 min). The nuclei-enriched fractions (5ml) were collected from the interphase between the Percoll® layers using a 5-ml pipette and the combined aliquots diluted in 23 ml gradient buffer (0.09 M hexylene glycol, 0.09 mM PIPES-KOH pH7, 1.83 mM MgCl2, 0.92 mM β-mercaptoethanol, 0.18% Triton X-100). To gently pellet the nuclei, the samples cushioned on 6 ml 30% Percoll® solution were centrifuged at 2,000 g and 4°C for 10 min. The isolated nuclei were resuspended in 1 ml sample buffer (20 mM TrisHCl pH7.4, 2 mM MgCl2, 150 mM NaCl, 5% glycerol, 5 mM DTT, cOmplete™ protease inhibitor (Roche)) and transferred into fresh 1.5 ml Eppendorf tubes and once washed in sample buffer (cenrifugation 1,000 g at 4°C for 15 min) and resupended in a final volume of 600µl sample buffer. Samples were treated with 1 µl DNase I (10 u/µl) and 2 µl of RNase A (10 mg/ml) for 15 min at 37°C and subsequently sonicated in a Bioruptor (Diagenode) water bath connected to a Minichiller cooling system (Huber) (6x 15 s “on”/15 s “off” at high intensity). After removal of debris (centrifugation at 16,000 g and 4°C for 15 min), supernatants were transferred into clean 2 ml Protein LoBind® tubes (Eppendorf). The protein concentration was determined by Bradford assay (Bradford, 1976) and equal amounts (i.e. 1 mg) were used for subsequent affinity purification. Immunoprecipitation was carried out with 25 µl GFP-trap Agarose beads (gta-20; Chromotek) in 2 ml sample buffer with Triton X-100 (0.1%) and EDTA (2 mM) after incubation at 4°C for 2.5 h at constant rotation (12 rpm). The protein-bound GFP-trap beads were washed four times with 300 µL of wash buffer (20 mM Tris-HCl pH7.4, 150 mM NaCl, 2 mM EDTA).

### Sample preparation and LC-MS/MS data acquisition

Proteins from GFP-trap enrichment were submitted to an on-bead digestion. In brief, dry beads were re-dissolved in 25 µL digestion buffer 1 (50 mM Tris, pH 7.5, 2M urea, 1mM DTT, 5 ng/µL trypsin) and incubated for 30 min at 30 °C in a Thermomixer with 400 rpm. Next, beads were pelleted, and the supernatant was transferred to a fresh tube. Digestion buffer 2 (50 mM Tris, pH 7.5, 2M urea, 5 mM CAA) was added to the beads; after mixing, the beads were pelleted, the supernatant was collected and combined with the previous one. The combined supernatants were then incubated o/n at 32 °C in a Thermomixer with 400 rpm; samples were protected from light during incubation. The digestion was stopped by adding 1 µL TFA and desalted with C18 Empore disk membranes according to the StageTip protocol [47]. Dried peptides were re-dissolved in 2% ACN, 0.1% TFA (10 µL) for analysis and measured without dilution. Samples were analyzed using an EASY-nLC 1200 (Thermo Fisher) coupled to a Q Exactive Plus mass spectrometer (Thermo Fisher). Peptides were separated on 16 cm frit-less silica emitters (New Objective, 75 µm inner diameter), packed in-house with reversed-phase ReproSil-Pur C18 AQ 1.9 µm resin (Dr. Maisch). Peptides were loaded on the column and eluted for 115 min using a segmented linear gradient of 5% to 95% solvent B (0 min : 5%B; 0-5 min -> 5%B; 5-65 min -> 20%B; 65-90 min ->35%B; 90-100 min -> 55%; 100-105 min ->95%, 105-115 min ->95%) (solvent A 0% ACN, 0.1% FA; solvent B 80% ACN, 0.1%FA) at a flow rate of 300 nL/min. Mass spectra were acquired in data-dependent acquisition mode with a TOP15 method. MS spectra were acquired in the Orbitrap analyzer with a mass range of 300–1750 m/z at a resolution of 70,000 FWHM and a target value of 3×10^6^ ions. Precursors were selected with an isolation window of 1.3 m/z. HCD fragmentation was performed at a normalized collision energy of 25. MS/MS spectra were acquired with a target value of 10^5^ ions at a resolution of 17,500 FWHM, a maximum injection time (max.) of 55 ms and a fixed first mass of m/z 100. Peptides with a charge of +1, greater than 6, or with unassigned charge state were excluded from fragmentation for MS^2^, dynamic exclusion for 30s prevented repeated selection of precursors.

### LC-MS/MS data data analysis

Raw data were processed using MaxQuant software (version 1.6.3.4, http://www.maxquant.org/) [48] with label-free quantification (LFQ) and iBAQ enabled [49]. MS/MS spectra were searched by the Andromeda search engine against a combined database containing the sequences from *A. thaliana* (TAIR10_pep_20101214; ftp://ftp.arabidopsis.org/home/tair/Proteins/TAIR10_protein_lists/) and sequences of 248 common contaminant proteins and decoy sequences. Trypsin specificity was required and a maximum of two missed cleavages allowed. Minimal peptide length was set to seven amino acids. Carbamidomethylation of cysteine residues was set as fixed, oxidation of methionine and protein N-terminal acetylation as variable modifications. Peptide-spectrum-matches and proteins were retained if they were below a false discovery rate of 1%.

Statistical analysis of the MaxLFQ values was carried out using Perseus (version 1.5.8.5, http://www.maxquant.org/). Quantified proteins were filtered for reverse hits and hits “identified by site” and MaxLFQ values were log2 transformed. Missing values were imputed from a normal distribution (1.8 downshift, separately for each column). After grouping samples by condition, only proteins with three valid values in at least one condition were retained for subsequent analysis. Statistically significant enrichment was performed by ANOVA followed by Honest True Difference (HSD) test for groups TRB1, TRB3, ESD1 with FDR<0.05.

### Protein expression and purification

A single colony of *E. coli* SoluBL21^TM^ (amsbio), carrying either *pET-28b-TRB1 or pET-28b-TRB3* was used to inoculate 5 ml preculture in LB-AMP (100 mg/ml ampicillin), and grown at 37 °C, 200 rpm overnight. The preculture was added to 1 l LB-AMP-medium and grown at 37 °C, 200 rpm until the OD_600_ was around 0.6 – 0.8. After addition of 1 mM IPTG the culture was transferred to 16 °C,200 rpm overnight. Bacterial cells were collected using a JLA 10.500 rotor (Beckman/centrifuge AvantiTM J-25) at 4000 rpm at 4 °C for 10 min. Afterwards, bacterial cells were resuspended in 40 ml ice-cold lysis buffer (50 mM NaPO_4_, 300 mM NaCl, 10 mM imidazole, 0.1 M PMSF at pH 7.5) and disrupted using sonication (Ultrasonic-Desintegrator, Branson) (Programme: Strength: 6, Duty cycle: 40, 3x 2 min). The cell debris was removed using a JA 25.50 rotor (Beckman/centrifuge AvantiTM J-25) at 13000 rpm, 4 °C for 30 min. For affinity purification, 500 µl of Ni-NTA Agarose beads (Qiagen) were washed three times with 5 ml lysis buffer, collected at 800 g,4 °C for 1 min and added to the cell lysate. After incubation at 4 °C, 12 rpm for 2 h then beads were collected, transferred to a fresh 5 ml Eppendorf tube and washed five times with 5 ml washing buffer (50 mM NaPO_4_, 300 mM NaCl, 20 mM imidazole, pH 7.5) at 4 °C, 12 rpm for 5 min. To elute proteins, the beads were incubated with 1.5 ml elution buffer (50 mM NaPO_4_, 300 mM NaCl, 250 mM imidazole, pH 7.5) at 4 °C, 12 rpm for 2 h. After collecting the beads at 4 °C, 800 g for 1 min, the supernatant was collected and dialysed in 500 ml dialysis buffer (50 mM NaPO_4_, 300 mM NaCl, pH 7.5) using Slide-A-Lyzer^TM^ Dialysis Cassettes (10K MWCO, 3 mL, ThermoFisher Scientific) to remove the imidazole. Dialysed proteins were collected in Protein LoBind Tubes (Eppendorf) and kept at 4 °C. Protein quantity was determined by Bio-Rad Protein Assay according to manufacturer’s instructions. To check protein integrity, 1 µg of protein was mixed 2x SDS-Loading buffer (126 mM Tris-HCl (pH 6.8), 20 % glycerol, 4 % SDS, 0.02 % bromophenol blue), incubated at 95 °C for 5 min, and run in 1x TGS buffer (Bio Rad) on a 1.5 mm, 12 % SDS-PAGE at 100 V for approximately 1.5 h. The SDS-PAGE was stained with Coomassie brilliant blue staining solution (1 g Coomassie Brilliant Blue (Bio-Rad), 500 ml MeOH, 100 ml glacial acetic acid, 400 ml H_2_O) and de-stained with H_2_O overnight.

### Microscale thermophoresis (MST)

Forward and reverse 5’-Cy3-labelled oligonucleotides of 28 bp were ordered from SIGMA-ALDRICH. Sequences originating from the *SEP3* promoter region were Cy3-*proSEP3*-telobox-Cy3: TTTAAATGT<u>TAGGGTTT</u>TTTGTAGGATT and Cy3-*proSEP3*-NonInter-Cy3: AAAAATATTTATATCACATCATTGTTAT). Two versions of the (C)RACCTA motif were Cy3-(C)AACCTAA-Cy3: CATCATGG<u>CAACCTAA</u>GGCTGGTACT AG and Cy3-(C)GACCTAA-Cy3: CATCATGG<u>CGACCTAA</u>GGCTGGTACTAG. A four-telobox-repeat (R4) oligomer Cy3-R4-telobox-Cy3: GGTTTAGGGTTTAGGGTTTAGGGTTTAG was published in [27]. Annealing was carried out in a heating block in dialysis buffer at a concentration of 10 µM sense and anti-sense oligonucleotides by first incubating at 95 °C for 15 min and subsequent slow cooling by switching off the heating block.

For all MST experiments, the Monolith NT.115 instrument (NanoTemper Technologies) and 1x dialysis buffer with Tween 20 (0.05 %) and BSA (1.25 mg/ml) were used. Oligomer fluorescence intensity, absorption, and bleaching was tested with the instrument’s green channel *via* the *Pretest* feature included in the machine’s MO.Control software (NanoTemper Technologies). Samples were prepared according to the suggested protocol included in the software. Oligomer concentration was adjusted to obtain ≥ 200 fluorescent counts at a laser power ≤ 80 %. These conditions were met at an oligomer concentration of 20 nM and an IR-laser power of 60 % for Cy3-*proSEP3*-telobox-Cy3 and Cy3-*proSEP3*-NonInter-Cy3 and of 80 % for Cy3-R4-telobox-Cy3, Cy3-(C)AACCTAA-Cy3, and Cy3-(C)GACCTAA-Cy3. Afterward, general TRB-telobox/telobox-like element interaction and suitability of different capillaries was tested *via* the Binding Check feature and samples were prepared as suggested by the software. For this purpose, the highest possible protein concentration was mixed with 20 nM of fluorescently labelled oligomers and incubated at RT and in the dark for 10 min. Afterward, the TRB-dsDNA mix was loaded onto Monolith NT.115 Premium Capillaries (NanoTemper Technologies) that prevented surface absorption, as TRB proteins tended to absorb to standard capillaries. Afterward, TRB-telobox/telobox-like element interaction was quantified by using the software’s *Binding Affinity* feature. For this MST assay, a dilution series was prepared according to the software’s instructions, using the beforehand determined laser powers and 20 nM of oligomer mixed with 8.5 µM to 260 pM of TRB protein. The TRB-DNA mix was incubated at RT in the dark for 10 min before being loaded onto Premium capillaries. Each measurement was repeated at least three times. Binding curves were analysed and K_D_ values were calculated with the MO.Affinity Analysis software (NanoTemper Technologies) according to the manufacturer’s instructions.

## Declarations

### Ethics approval and consent to participate

Not applicable

### Consent for publication

Not applicable

### Availability of data and materials

The datasets supporting the conclusions of this article are available in the European Nucleotide Archive (ENA), under the project accession number PRJEB63124.

### Authors’ contributions

F.T., K.K., M.M. and S.Z. conceived the study. K.K., M.M., S.Z., P.T. conducted the experiments. S.S. and H.N. set-up and carried out MS-MS analysis. M.M, K.K., P.S. and F.T. analyzed the results. F.T., K.K. and M.M wrote the manuscript. All authors reviewed the manuscript.

## Supplemental Information

### Supplemental Figures

Supplemental Figure 1. Expression of flowering time pathway genes in *trb* mutants

Supplemental Figure 2. Bioinformatics pipeline for ChIP-seq analysis and ChIP-seq quality control

Supplemental Figure 3. Characterization of TRB1-3 ChIP-seq data

Supplemental Figure 4. Pearson’s correlation matrix of enriched gene ontology (GO) terms across all combinations of TRBs and complex categories

**Supplemental File 1: Details of RNAseq analysis**

Tab sig_1_wt_inS: DEGs trb1-2 vs Col-0

Tab sig_2_wt_inS: DEGs trb2-2 vs Col-0

Tab sig_2_wt_inS: DEGs trb3-2 vs Col-0

Tab sig_12_wt_inS: DEGs trb1-2 trb2-3 vs Col-0

Tab sig_13_wt_inS: DEGs trb1-2 trb3-2 vs Col-0

Tab sig_23_wt_inS: DEGs trb2-3 trb3-2 vs Col-0

Tab vst countmatrix: vst countmatrix across all genotypes for all DEGs

Tab PAM countmatrix: row scaled countmatrix across all genotypes for all DEGs with PAMclusters indicated

Tab detected genes: all genes with acceptable read counts (total >50 across all samples) in RNAseq

Tab GO PAM: GO-term enrichment for DEGs per PAM cluster

**Supplemental File 2: Details of ChIP-Seq analysis**

Tab peaks_TRB1: Annotated ChIP-Seq Peaks of TRB1:YFP

Tab peaks_TRB2: Annotated ChIP-Seq Peaks of TRB2:YFP

Tab peaks_TRB3: Annotated ChIP-Seq Peaks of TRB3:YFP

**Supplemental File 3: Details of genomic binding analysis**

Tab bins: Table of 11 regulatory protein occupancy of 500 bp wide genomic Bins, assigned regulatory complexes, and combination of TRB paralogs

Tab genes: Table of 11 regulatory protein occupancy of genes, associated RNAseq Cluster, assigned regulatory complexes, and combination of TRB paralogs

**Supplemental File 4: Details of IP-MS-MS analysis**

Tab raw: MaxQuant result

Tab ANOVA_HSD_full: log2 transformed, filtered and imputed LFQ values, result of ANOVA and HSD annotated to all protein

Tab: Significant: All proteins part of significant HSD groups

**Supplemental File 5: Details of DNA-Motif analysis**

Supplemental_File_5.zip: FASTA sequences used as input for the XSTREME analysis of the MEME-Pipeline, as well as directories containing the outputs

**Supplemental File 6: Details of GO-Term enrichment analysis**

Tab All_GO_Terms: Table containing all enriched (p<=0.05) GO-terms, their associated complexes, and co-bound TRB-paralogs

