## Supplemental_Figures_and_Tables for "Epigenetic gene regulation is controlled by distinct regulatory complexes utilizing specialized paralogs of TELOMERE REPEAT BINDING FACTORS"

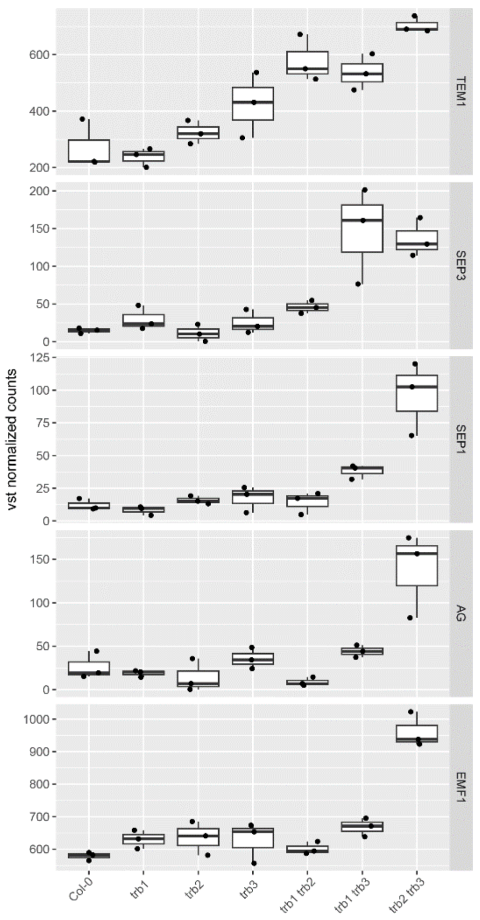


**Supplemental Figure 1. Expression of flowering time pathway genes in *trb* mutants**. Boxplots show variance stabilized read counts of RNA-seq analysis (n= 3 biological and experimental replicates) in trb single and double mutants. All genes were part of transcriptional cluster 4 as described in Figure 1. Boxplots show innerquartile range and median as line, each data indicated as dot.

A

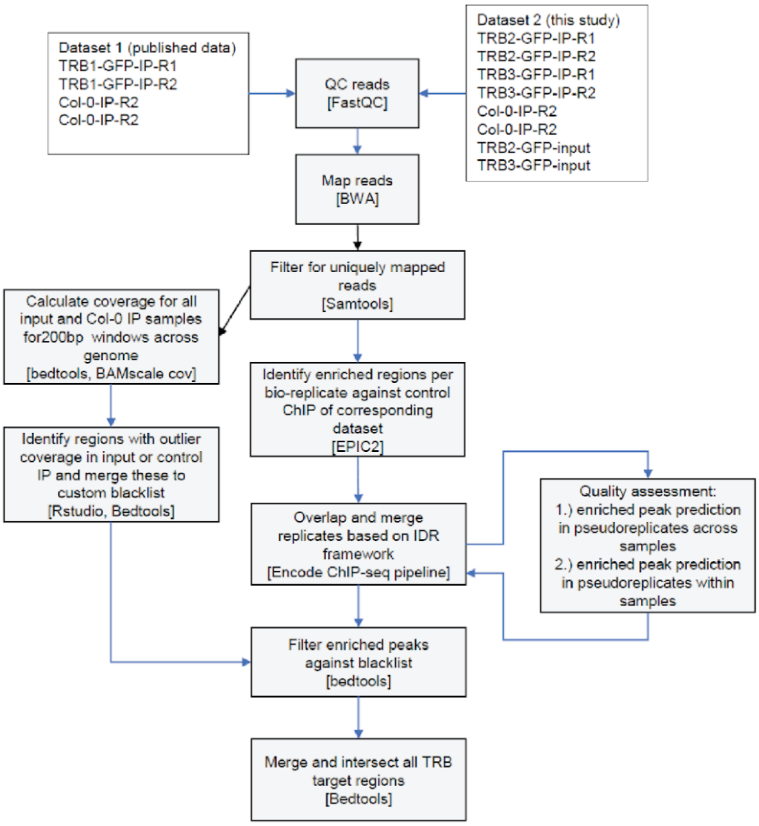


B

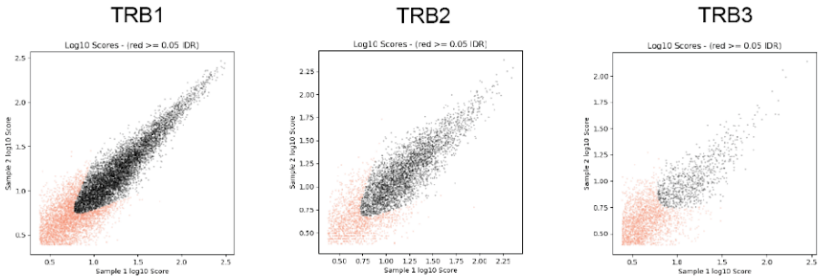


C

| TRB1 | TRB2 | TRB3 | IDR type |
|---|---|---|---|
| 7483 | 3771 | 845 | true replicates |
| 8690 | 4321 | 1450 | pseudo-replicates across samples |
| 6607 | 3064 | 1155 | pseudo-replicates within replicate 1 |
| 5545 | 3119 | 613 | pseudo-replicates within replicate 2 |
| 1.2 (pass) | 1.1 (pass) | 1.7 (pass) | Np/Nt |
| 1.2 (pass) | 1.0 (pass) | 1.9 (pass) | N1/N2 |

**Supplemental Figure 2. Bioinformatics pipeline for ChIP-seq analysis and ChIP-seq quality control. A)** Flowchart of the pipeline used for identification of enriched regions by ChIP-seq. **B)** Comparison of sample scores between 2 replicates (biological and experimental) using the Irreproducible Discovery Rate (IDR) framework as implemented by Encode. Samples were ranked by score determined by EPIC2, values with an IDR < 0.05 were considered as positive) **C)** Number of peaks identified by the pipeline in true replicates, pseudo-replicates and within sample replicates as indicated by IDRtype.

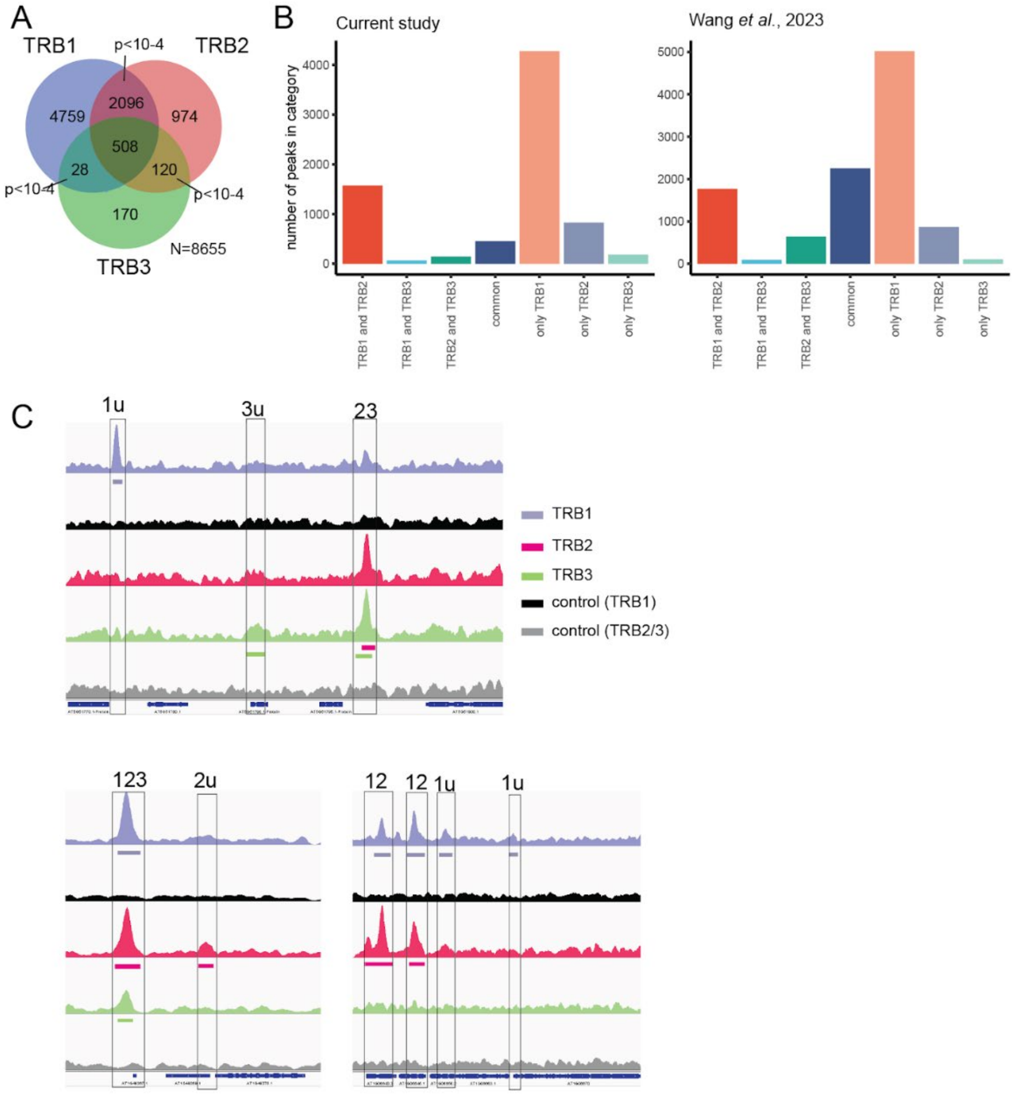


**Supplemental Figure 3. Characterization of TRB1-3 ChIP-seq data. A)** Venn diagram of peaks for TRB1-3. Peaks were backannotated after merging all peaks. Statistical testing by permutation as implemented in Genome Association Tester (GAT). **B)** Comparison of TRB1-3 peak categories identified in this study (left panel) and in a previous study by Wang et al. 2023. C) Genome browser tracks showing typical peaks (1u, 2u, 3u: only detected in TRB1, TRB2,TRB2. 12, 23: Peaks detected for TRB1 and TRB2 or TRB2 and TRB3. 123: peaks commonly detected). Background (control) shows ChIP-seq libraries prepared from Col-0. All tracks are normalized to Counts per million mapped fragments (CPM).

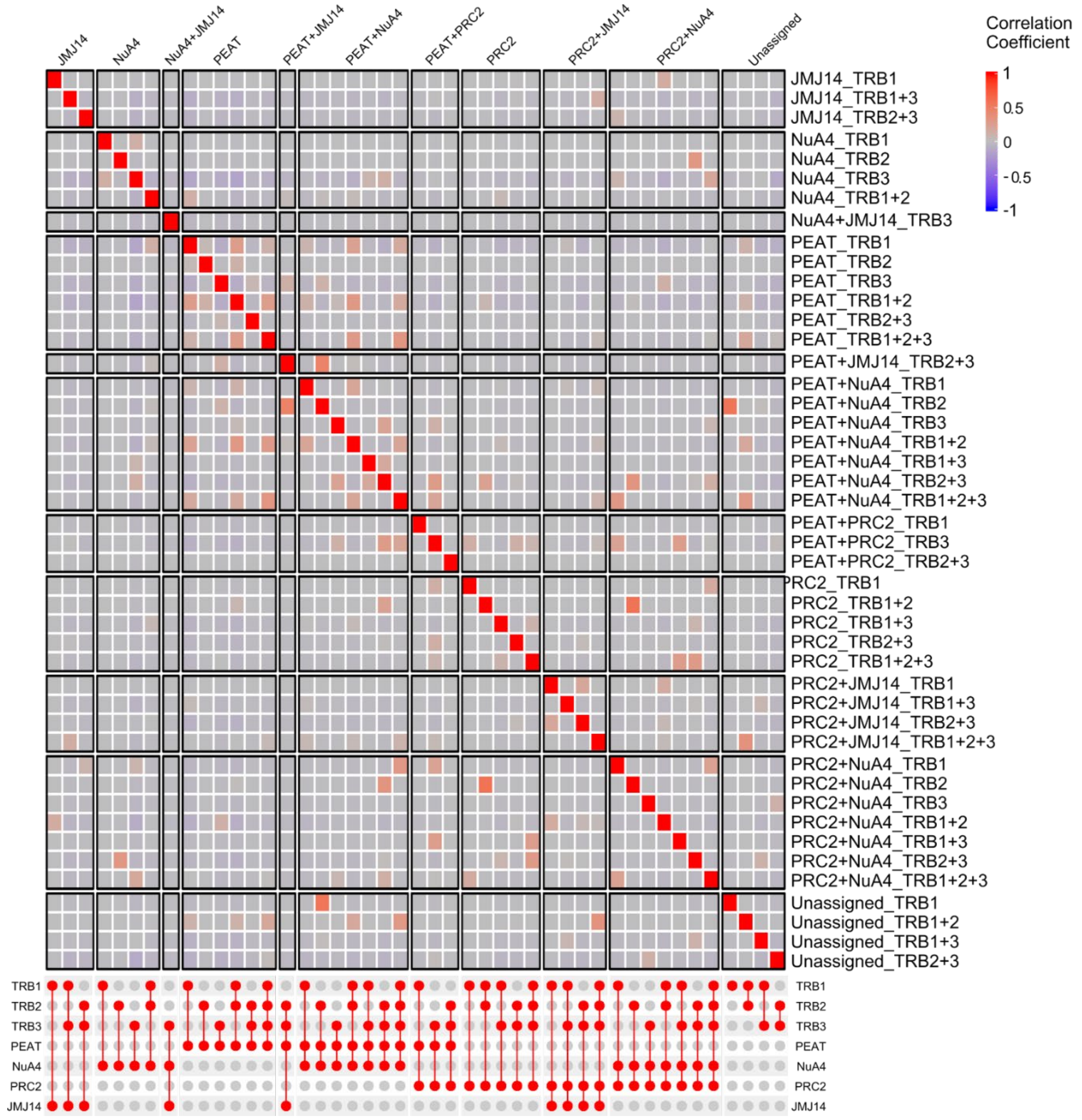


**Supplemental Figure 4:** Pearson's correlation matrix of enriched gene ontology (GO) terms across all combinations of TRBs and complex categories. Combinations that do not enrich any GO-Terms have been excluded.
